# The Segmentome: a novel representation of the DNA methylome identified by a Bayesian HMM framework

**DOI:** 10.64898/2026.11.01.763464

**Authors:** Mahdi Anvari, Niloofar Torki, Mahya Mehrmohamadi

## Abstract

**Motivation:** DNA methylation is organized along the genome into regions of relatively stable methylation (i.e. fully methylated or fully unmethylated regions), separated by transition zones. Differences in the boundaries of these methylation zones across cells can result from various regulatory and epigenetic mechanisms. Most DNA methylome studies, however, are focused on the detection of differentially methylated regions (DMRs), and this organization within a single methylome remains under explored. Furthermore it is unclear whether a segment-level descriptor can capture cell identity.

**Results:** We introduce the segmentome, a segment-level representation of the methylome, and an unsupervised Bayesian hidden Markov model that infers it from WGBS data without predefined methylation thresholds or labels. The model combines CpG methylation levels with read-pattern entropy, local CpG correlation and CpG count to divide the genome into unmethylated, methylated and transition zones, the last characterized by heterogeneous read patterns. Applied to 207 samples of 39 human cell types, segmentome profiles grouped samples by cell type in an unsupervised manner (adjusted Rand index 0.55, adjusted mutual information 0.72). We next generated a segmentome atlas consisting of top segmentome biomarkers per human cell-type. These markers were capable of cell-of-origin determining in cell-free DNA and white blood cell samples. These results suggest that segmentation is a unique framework for analyzing DNA methylome.

## 1 Introduction

DNA methylation is a key epigenetic mechanism that plays role in gene expression regulation and maintenance of genome integrity and cellular identity (Bogdanović & Lister, 2017). In mammals DNA methylation occurs mainly at CpG sites, and its distribution varies substantially across genomic regions and cell types (Smith & Meissner, 2013). These patterns are not random: neighboring CpGs often show coordinated methylation states, which results in contiguous genomic regions with relatively stable methylation, separated by transition points known as methylation boundaries (Naumann et al., 2009; Sofer et al., 2013; Su et al., 2013). Characterization of this spatial organization is therefore part of understanding how methylation contributes to genome regulation.

Methylation boundaries may carry regulatory information because the variable fraction of the methylome is concentrated at regulatory elements. Most CpGs are similarly methylated across cell types; the CpGs that differ lie mainly at distal regulatory elements such as enhancers, where methylation varies with transcription factor binding and chromatin state (Stadler et al., 2011; Ziller et al., 2013; Kundaje et al., 2015). Transitions between stable methylation states may therefore coincide with sites of regulator binding and chromatin change. If so, their positions would be expected to differ between cell types, developmental stages and disease states. Cancer offers one example: methylation becomes more variable within large epigenetic domains (Hansen et al., 2011). Mapping the positions of boundaries, and how they shift between conditions, could therefore reveal regulatory differences that methylation levels at individual CpGs or predefined regions do not capture.

Technologies that measure DNA methylation at single CpG resolution and fragment-level are ideal for identifying the location of boundaries where DNA methylation states change across the genome. For instance whole-genome bisulfite sequencing (WGBS) measures methylation at single CpG resolution across the genome, without restricting the analysis to predefined genomic regions (Lister et al., 2009; Laird, 2010). Each sequenced fragment also retains the methylation pattern of several consecutive CpGs, which shows how neighboring CpGs vary together on the same molecule (Guo et al., 2017).

Various computational strategies have been developed to analyze WGBS data and characterize DNA methylation states genomewide (Wreczycka et al., 2017). The major focus of these studies has been the identification of differentially methylated regions (DMRs) between cases and controls in various contexts. DMR analysis generally has two steps: differentially methylated cytosines (DMCs) are identified by comparing methylation levels at individual CpGs between predefined groups with statistical tests, and neighboring DMCs are then grouped into contiguous DMRs by distance criteria (Piao et al., 2021; Shafi et al., 2018). DMRs are therefore defined by differences in methylation between groups, not by the organization of methylation states along a single genome-wide profile (Chen et al., 2016).

Previous attempts at describing the organization of DNA methylation across one methylome profile have certain limitations that motivated us to undertake the present study. MethylSeekR, for example, identifies partially methylated domains and regions of reduced methylation from CpG methylation fractions (Burger et al., 2013). Hidden Markov models (HMMs) have been used to represent methylation states while accounting for the dependence between neighboring genomic positions (Malonzo & Lähdesmäki, 2023), although many HMM-based methods were developed mainly to detect differential methylation (Ji, 2019; Shen et al., 2017). An unsupervised, comprehensive characterization of the DNA methylome across human cell types that represents the transitions between stable methylation regions explicitly is currently missing.

We therefore developed an unsupervised Bayesian HMM that infers the methylation organization of a single methylome from WGBS data (Figure 1B–D). A sliding window across the genome quantifies methylation, followed by merging of adjacent windows in the same state into segments, such that boundaries are the points at which the inferred state changes. We refer to the resulting set of segments as the *segmentome*. Unlike a methylome profile, which lists methylation measurements at individual CpG sites, the segmentome summarized the methylome into spatially organized genomic regions with inferred methylation states. We present the segmentome atlas across 39 human cell types and identify cell-type-specific-segments.

**Figure 1.**
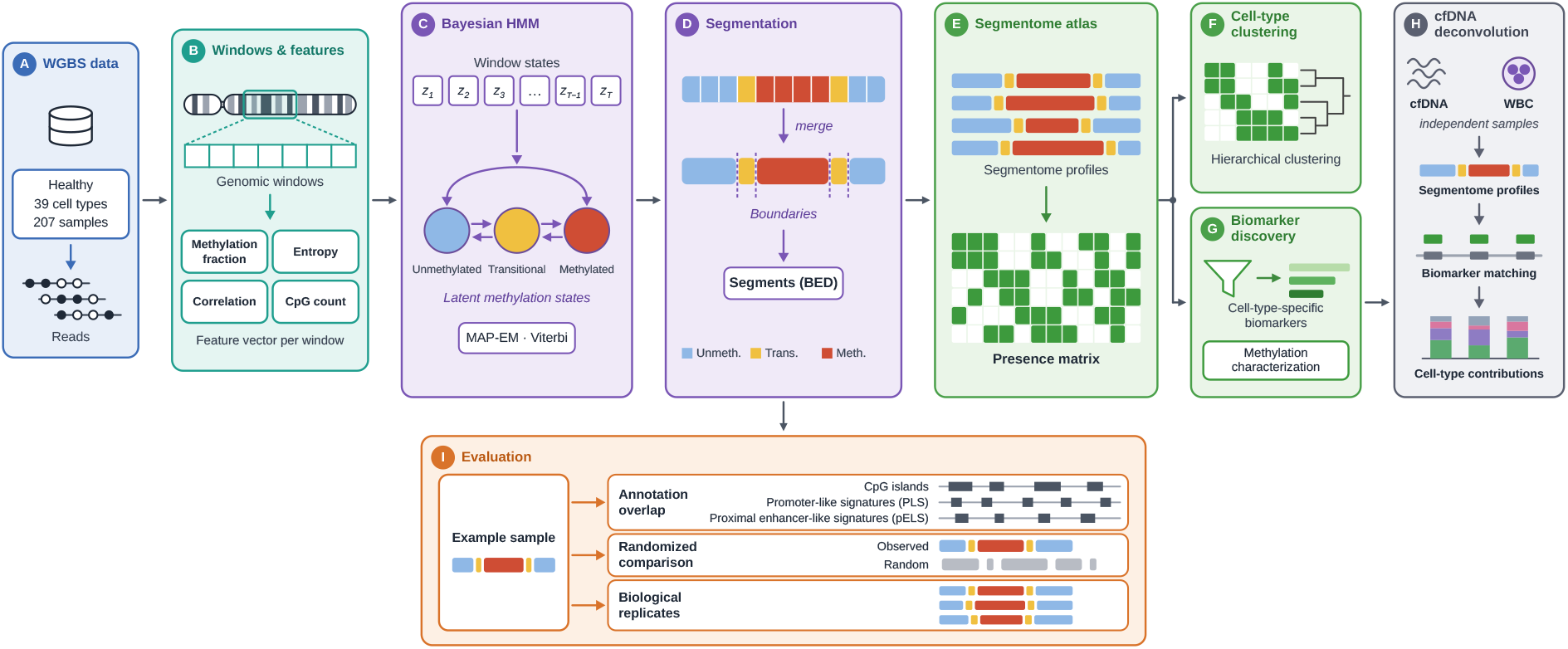
Overview of the segmentome discovery framework. (A) WGBS data were obtained from a public atlas of healthy human cell types (39 cell types, 207 samples; (Loyfer et al., 2023)). Each read carries the methylation status of its CpG sites (filled, methylated; open, unmethylated). (B) The genome is divided into consecutive 50-bp windows, and reads are assigned to the windows they overlap. Each window is summarized by four features: methylation fraction, read-pattern entropy, local CpG correlation and CpG count. (C) For each chromosome, the ordered window-level feature vectors are modeled by a Bayesian hidden Markov model (HMM) with three latent states (unmethylated, transition and methylated). Parameters are estimated by maximum a posteriori expectation–maximization (MAP-EM), and the most probable state sequence (z1…zT) is decoded with the Viterbi algorithm. (D) Adjacent windows in the same state are merged into segments, which are exported as BED intervals. (E) The segmentation is applied to every sample, and the resulting segmentome profiles are combined into a binary presence matrix recording which segments are detected in each sample. (F) Hierarchical clustering of the presence matrix tests whether profiles group samples by cell type. (G) Cell-type-specific biomarker segments are selected from the presence matrix, and their methylation levels are compared between target and background samples. (H) Segmentome profiles of independent cell-free DNA (cfDNA) and white blood cell (WBC) samples are matched to the biomarker sets to estimate cell-type contributions. (I) Evaluation of the segmentation on a representative oligodendrocyte sample (GSM5652222): overlap of its segments with genomic annotations (CpG islands, promoter-like signatures [PLS] and proximal enhancer-like signatures [pELS]), comparison with randomized segments, and reproducibility in three biological replicates (GSM5652221, GSM5652220 and GSM5652219).

We then evaluate the segmentome as a representation for comparing methylomes across cell types, showing potential application in liquid biopsy tissue-of-origin determination.

## 2 Materials and methods

### 2.1 Study design

We developed a Bayesian segmentation framework that identifies genomic regions with locally stable DNA methylation, and the transitions between them, from whole-genome bisulfite sequencing (WGBS) data of a single methylome (Figure 1). The study had three parts. First, the framework was built and evaluated on a representative oligodendrocyte sample: CpG-level methylation profiles were prepared, the genome was divided into windows described by four features, a hidden Markov model (HMM) assigned a methylation state to each window, and adjacent windows in the same state were merged into segments (Sections 2.2–2.6). Second, the segments were validated by their overlap with annotated genomic regions and by comparison with randomized segmentations, and their reproducibility was tested in three biological replicates (Sections 2.7–2.9). Third, the framework was applied to a DNA methylation atlas of human cell types, and the resulting segmentome profiles were used for clustering, biomarker discovery and the analysis of cell-free DNA (Sections 2.10–2.13).

The segmentation is unsupervised. No genomic region was assigned to a predefined methylation class during model fitting; the HMM inferred latent states from the window-level features alone, and the inferred states were characterized afterwards by their methylation profiles. This allows the model to identify both relatively stable methylation regions and the intervals in which methylation changes.

### 2.2 WGBS dataset

We used publicly available WGBS data from the human DNA methylation atlas of Loyfer et al. (2023). The atlas contains deeply sequenced methylomes of healthy human cell types, which were purified by fluorescence-activated cell sorting (FACS) and sequenced to a mean depth of at least 30×. We analyzed all 207 samples of the atlas available under GEO accession GSE186458, which represent 39 cell types. The data were provided in PAT format, which preserves the methylation pattern of the consecutive CpGs covered by each sequencing fragment. PAT files were converted to CpG-level methylation data with wgbstools (Loyfer et al., 2026), which retains the genomic position and methylation status of each CpG.

A representative oligodendrocyte sample (GSM5652222) was used to develop and evaluate the segmentation procedure. Once the workflow was fixed, the same procedure was applied to all other samples to build the segmentome atlas.

The analysis was restricted to the autosomes (chromosomes 1– 22) and, within each chromosome, to the WGBS-callable region. We defined this region as the interval from the first to the last CpG site of the chromosome in the wgbstools hg19 CpG index (the reference file CpG.bed.gz that wgbstools builds from the hg19 genome sequence with the init_genome command), that is, the part of the chromosome in which WGBS reads can cover CpGs. The callable regions cover 2,796,121,515 bp, 97.1% of the 2,881,033,286 bp of the hg19 autosomes. They are shorter than the chromosomes mainly on the acrocentric chromosomes 13, 14, 15, 21 and 22, which begin with 9–20 Mb without sequence in hg19 (Supplementary Figure S1A). Because each region is a single interval per chromosome, it also contains internal gaps of the assembly. Each chromosome was processed independently so that no artificial transitions arise between chromosomes. All coordinates follow the hg19 assembly of the PAT files, which was used consistently for CpG indexing, genomic coordinates and annotation.

### 2.3 Genomic window construction

Methylation at nearby CpGs is correlated, and the correlation decreases with distance (Guo et al., 2017; Lövkvist et al., 2016; Nautiyal et al., 2010). WGBS fragments also retain the methylation pattern of several consecutive CpGs, which a single CpG does not show. To use this local information, each chromosome was divided into consecutive, non-overlapping windows of 50 bp, and the genomic order of the windows was kept throughout the analysis. A fragment was assigned to every window with which it shared at least 1 bp. When a fragment spanned the boundary between windows, it was split, so that each window received only the part of the fragment within its coordinates.

### 2.4 Feature extraction

Four features were calculated for each window from the CpG observations it contains: methylation fraction, read-pattern entropy, local CpG correlation and the number of CpGs. Windows with fewer than two CpGs (the minimum for the local CpG correlation) or without reads were excluded. The four features of a window form the observation vector of the HMM.

#### 2.4.1 Methylation fraction

The methylation fraction is the proportion of methylated CpG observations among all CpG observations in the window.

### 2.4.2 Read-pattern entropy

Each fragment covering CpGs in the window was represented by its methylation pattern, that is, the methylation states of the CpGs it covers in the window, and fragments with the same pattern formed a pattern class. The read-pattern entropy is the Shannon entropy of the distribution of pattern classes,

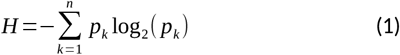

where 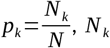, *N* _*k*_ is the number of fragments in pattern class k, and N is the number of fragments in the window. Higher entropy indicates a more diverse set of methylation patterns among fragments, and lower entropy indicates that a few patterns account for most of them.

#### 2.4.3 Local CpG correlation

The local CpG correlation is the proportion of adjacent CpG pairs with the same methylation state among all adjacent CpG pairs on the fragments covering the window (a fragment with L CpGs contributes L − 1 pairs). It measures how consistently neighboring CpGs behave together within fragments. A value of 1 means that all adjacent pairs are concordant.

#### 2.4.4 Number of CpGs

The number of CpGs is the number of distinct CpG sites represented in the window. It reflects the CpG density of the window and the amount of CpG-level information available for it.

### 2.5 Bayesian hidden Markov model

For each sample and autosome, the ordered sequence of window feature vectors was modeled with a Bayesian HMM in which every window belongs to one of three latent states. State-transition probabilities capture the dependence between neighboring windows, so that the state of one window informs the states of its neighbors. Given the state, the four features were modeled as independent, each with its own emission distribution: a zero-andone-inflated Beta distribution for the methylation fraction and for the local CpG correlation, a zero-inflated Gamma distribution for the read-pattern entropy, and a Poisson distribution for the number of CpGs. The emission likelihood of a window is the product of the four feature likelihoods.

The parameters were estimated by maximum a posteriori (MAP) inference with an expectation–maximization (EM) algorithm. In the E step, the forward–backward algorithm gave the posterior state probabilities and the posterior probabilities of transitions between neighboring states. In the M step, the transition probabilities and the emission parameters were updated using these posteriors as weights. Iteration stopped when the relative improvement of the log-likelihood fell below a tolerance or when the maximum number of iterations was reached. The most probable state sequence was then obtained with the Viterbi algorithm. The states were ordered by their mean methylation fraction and labeled unmethylated (lowest), transition (intermediate) and methylated (highest).

### 2.6 Segmentation

Adjacent windows assigned to the same state were merged into continuous segments when the gap between them did not exceed 1,000 bp. Gaps arise where windows were excluded by the filters in Section 2.4. A segment spans from the start of its first window to the end of its last window, including any gaps. Windows were merged only within a chromosome. The segmentation of each sample was exported in BED format with the coordinates and state of each segment and the sample identifier.

### 2.7 Randomized segmentations

To test whether the genomic distribution of the segments differs from chance, randomized segmentations were generated for each chromosome within the WGBS-callable space. The lengths of all observed segments on the chromosome were kept and their order was randomly permuted. The callable space not occupied by segments was then distributed randomly among the N + 1 gaps before, between and after the N randomized segments, with gap sizes drawn so that they sum exactly to the free space. The randomized segments therefore do not overlap each other, stay within the callable region, and keep the number and length distribution of the observed segments on each chromosome, while their positions are random. The procedure was repeated 1,000 times.

### 2.8 Overlap analysis and statistical validation

#### 2.8.1 Genomic annotations and overlap

Segments were compared with three sets of annotations in hg19 coordinates. CpG islands (n = 26,641) were taken from the cpgIslandExt track of the UCSC Genome Browser (Kent et al., 2002). Promoter-like signatures (PLS; n = 45,890) and proximal enhancerlike signatures (pELS; n = 242,790) were taken from Registry V4 of candidate cis-regulatory elements (cCREs) of the ENCODE project (Moore et al., 2026) downloaded from the SCREEN portal (https://screen.wenglab.org/downloads); these classes were first defined in the earlier registry (Moore et al., 2020). The registry is released in GRCh38 coordinates, and we used the hg19 file provided by the portal (cCRE.hg19.bed), in which the coordinates were lifted from GRCh38 to hg19. The file lists each element with its class; the elements of the classes PLS and pELS were selected, and the other classes of the registry were not used. Segments and annotations were intersected with bedtools intersect (Quinlan & Hall, 2010). When an annotated element overlapped several segments, the segment with the largest overlap was kept as its best-overlapping segment. The same procedure was applied to the randomized segmentations.

#### 2.8.2 Overlap metrics

Six metrics were calculated. Global coverage is the total overlap divided by the total length of the annotated elements. The global Jaccard index is the total overlap divided by the length of the union of segments and annotated elements. Feature recall is the fraction of elements overlapped by at least one segment. For each element and its best-overlapping segment, coverage is the overlap divided by the element length, precision is the overlap divided by the segment length, and the Jaccard index is the overlap divided by the union of the two; their averages over all elements are the mean coverage, mean precision and mean Jaccard index.

#### 2.8.3 Empirical significance and fold change

Each metric was calculated for the observed segmentation and for each of the 1,000 randomized segmentations, which gave a null distribution for each metric. The one-sided empirical P value was

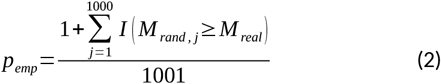

where *M*_*real*_ is the metric of the observed segmentation, *M* _*r and, j*_is the metric of randomized segmentation j, and I is the indicator function. Adding one to the numerator and the denominator avoids a P value of zero, and the smallest possible value is 1/1001, about 0.001. The size of the association was expressed as the fold change, the observed metric divided by the mean of the metric over the randomized segmentations; a fold change of 1 is the random expectation. In addition, the per-element coverage distributions of the observed segmentation and of one randomized segmentation were compared with two-sample Kolmogorov–Smirnov and two-sided Mann–Whitney U tests.

### 2.9 Reproducibility across biological replicates

To test whether the segmentation is reproducible, the same work-flow (Sections 2.3–2.8) was applied independently to three further oligodendrocyte WGBS samples (GSM5652221, GSM5652220 and GSM5652219) and compared with the initial sample GSM5652222.

At the window level, windows were matched by chromosome, start and end coordinates, and the overlap of the window sets and of the windows with the same state was shown as Venn diagrams. For each pair of samples, the agreement of state assignments was summarized as the proportion of windows of each state in one sample that received each state in the other. At the segment level, for each segment of the initial sample, the segment with the largest overlap in each replicate was identified and the pair was classified as the same state, adjacent states (transition and a stable state), opposite states (methylated and unmethylated) or no overlap. The overlap metrics of Section 2.8 were calculated for each sample and compared.

### 2.10 Segmentome atlas

The segmentation was applied to all 207 samples of the atlas (39 cell types) with the same configuration, so that the profiles arecomparable. The set of segments of a sample is its segmentome profile, and a segment is identified by its chromosome, start position and end position. The union of the segments of all samples formed the segment pool, which served as a common feature space. A binary presence matrix, with samples as rows and segments as columns, recorded whether each segment was present in the segmentome profile of each sample.

To characterize the growth of the segment pool, the samples were added one at a time in alphabetical order of their names, and the cumulative number of unique segments was recorded af-ter each sample. The order was fixed and was not randomized, so the number of new segments contributed by a sample depends on the samples added before it. A plateau of this curve would indicate that additional samples contribute progressively fewer segments that were not observed before.

#### 2.10.1 Representation relationships and correction of the presence matrix

Segments that differ only slightly in their boundaries count as separate features, which makes the presence matrix redundant and may affect clustering and biomarker discovery. We therefore defined two segments as representing each other when the length of their overlap was at least 90% of the length of the longer segment. The relationship was not assumed to be transitive: a relationship between segments A and B and one between B and C does not imply one between A and C. The corrected presence matrix was calculated from the initial presence matrix: when one segment of a related pair was present in a sample, its counterpart was also recorded as present.

### 2.11 Cell-type clustering

The ability of segmentome profiles to distinguish samples by cell type was evaluated using agglomerative hierarchical clustering based on the corrected binary presence matrix. To reduce the influence of uninformative features, segments were ranked according to their variance across samples, and the 28,000 most variable segments were selected to construct a reduced presence matrix. UMAP (McInnes et al., 2018) was applied to this matrix to transform the selected features into a 164-dimensional embedding of the samples (number of neighbors 30, minimum distance 0.1, Euclidean metric). Agglomerative hierarchical clustering was performed on this representation using Euclidean distance and complete linkage, which defines the distance between two clusters as the maximum pairwise distance between their constituent samples.

The resulting dendrogram was cut into 39 clusters, the number of annotated cell types, and compared with the original cell-type annotations using the adjusted Rand index (ARI), the adjusted mutual information (AMI) and purity. ARI and AMI measure the agreement between two partitions and are corrected for chance (1 for identical partitions, about 0 for random ones). Purity is the fraction of samples that belong to a cluster whose most frequent cell type is their own. Cell-type annotations were excluded from feature selection, dimensionality reduction and clustering and were used only for this evaluation. This approach enabled assessment of the discriminatory information captured by the segmentome representation in an unsupervised setting.

### 2.12 Biomarker discovery

#### 2.12.1 Segmentome cell-type-specific biomarkers

Cell-type-specific biomarkers were identified with the corrected presence matrix. For each cell type, a segment was a candidate biomarker if it met three criteria: its target prevalence, the proportion of samples of the target cell type in which the segment was present, was at least 70%; its background prevalence, the proportion of samples of all other cell types in which it was present, was below 2%; and its prevalence in any single non-target cell type was no greater than 20%. Candidates were ranked within each cell type by their target prevalence, from highest to lowest; candidates with equal target prevalence were ranked by background prevalence, from lowest to highest, then by the highest prevalence in any single non-target cell type, from lowest to highest, and any remaining ties were broken at random. The top 25 or the top 250 candidates per cell type were retained (all candidates if fewer were available; endothelial cells had only 18 candidates, so all 18 were retained). The top 25 were used to assess how well the biomarkers separate cell types: Euclidean distances between samples were calculated from the presence of these biomarkers and averaged for each pair of cell types. The top 250 were used for the analysis of methylation characteristics (Section 2.12.2).

#### 2.12.2 Methylation characteristics of segmentome biomarkers

To test whether segmentome biomarkers carry information that differs from that of conventional methylation biomarkers, we calculated their methylation levels in the underlying WGBS data. For each cell type, the mean methylation β value of each segment was calculated across the samples of the target cell type (target β) and across the samples of all other cell types (background β), and the fold change was the target β divided by the background β. The same procedure was applied to three sets: the segmentome biomarkers, the methylation biomarkers, and randomized control segments. The controls were generated by shuffling the segmentome biomarker intervals within the WGBS-callable regions, and randomized segments without sequencing reads were excluded. For each set, the fold changes were transformed with log_2_(fold change + 1), and the distributions of the three sets were compared pairwise with two-sample Kolmogorov–Smirnov tests, whose statistic is the maximum absolute difference between the two empirical cumulative distribution functions. The P values were adjusted for multiple comparisons with the Holm procedure.

### 2.13 Cell-free DNA deconvolution

We evaluated the segmentome biomarkers on 23 paired cell-free DNA (cfDNA) and white blood cell (WBC) samples from 23 healthy donors (Loyfer et al., 2023; GEO series GSE186458, samples GSM6810003–GSM6810025 for cfDNA and GSM6810026–GSM6810048 for WBC). These samples were not among the 207 atlas samples that were used to define the biomarkers. Segmentome profiles were generated for all 46 samples with the same trained model and segmentation procedure (Sections 2.5–2.6). For each sample, the number of detected biomarkers of each cell type was determined by matching its profile against the corresponding cell-type-specific biomarker set (a biomarker was counted as detected when a segment with identical chromosome, start and end coordinates was present in the profile of the sample), which contained all biomarkers identified for that cell type, not only the topranked ones. This gave a sample-by-cell-type matrix of raw counts (C).

To account for the different sizes of the biomarker sets, the counts were divided by the number of biomarkers in the set of the cell type (N = C/M, where M is the set size). The resulting values were then normalized across cell types within each sample by dividing each value by the sum over all cell types in that sample (R = N/ΣN), so that the values of each sample sum to one. The normalized profiles describe the relative representation of cell-type-specific biomarkers in cfDNA and WBC samples. They are comparative signatures and not calibrated estimates of cell-type proportions.

## 3 Results

### 3.1 Development of a computational framework for segmentation of the DNA methylome

In order to understand and characterize methylation state boundaries across the genome, we obtained publicly available WGBS data from the human DNA methylation atlas (Loyfer et al., 2023). We started with a single sample initially (oligodendrocyte sample GSM5652222), for the development and evaluation of our method (Figure 1B–D). The genome was divided into consecutive, non-overlapping 50-bp windows. Windows with fewer than two CpG sites or without reads were excluded, and the analysis was restricted to regions callable in the WGBS data (Supplementary Figure S1A). In total, 5,381,758 autosomal windows were retained for modeling, and the HMM assigned each of them to an unmethylated, transition or methylated state (see Methods for more details).

Methylated windows were the most frequent (62.1%), followed by transition (28.7%) and unmethylated windows (9.2%) (Supplementary Figure S1B). The number of windows per chromosome varied with chromosome size and with the amount of callable sequence, which is reduced on the acrocentric chromosomes 13, 14, 15, 21 and 22 (Supplementary Figure S1C).

Each window was then described by four features: methylation fraction (beta-value), read-pattern entropy, local CpG correlation, and total CpG count. Next, each window was assigned to one of three states, unmethylated, transition or methylated, which are inferred from these features without predefined methylation thresholds or biological labels. As expected, the four features differed among the states (Supplementary Figure S2A; values are means across chromosomes). Methylation fraction was lowest in unmethylated windows (0.021), highest in methylated windows (0.963) and intermediate in transition windows (0.701). Transition windows had the highest read-pattern entropy (2.109, versus 1.473 for unmethylated and 1.232 for methylated windows) and the lowest local CpG correlation (0.667, versus 0.959 and 0.928), indicating more diverse read-level methylation patterns and weaker agreement between neighboring CpGs than in either sta-ble state. Unmethylated windows had the highest CpG count (4.245, versus 2.524 for transition and 2.450 for methylated windows). Because the mean methylation fraction of transition windows was closer to that of methylated windows, the transition state is defined mainly by heterogeneous read patterns rather than by an intermediate methylation level alone. The state profiles were similar across chromosomes: in a principal component analysis of the chromosome-specific profiles, the 22 autosomes grouped by state rather than by chromosome (Supplementary Figure S2B).

The mean transition probabilities across chromosomes are shown in Supplementary Figure S2C. The unmethylated and methylated states had high self-transition probabilities (0.870 and 0.791), so they tended to persist across consecutive windows, whereas the transition state had a lower self-transition probability (0.542). Changes out of the transition state were asymmetric, going to the methylated state (0.427) far more often than to the unmethylated state (0.031). Direct switches between the two stable states were rare (0.029 from unmethylated to methylated and 0.004 in the opposite direction); the stable states switched mainly into the transition state (0.102 and 0.205, respectively). These probabilities varied little among chromosomes (standard deviation ≤ 0.008 for the stable states and ≤ 0.049 for the transition state; Supplementary Figure S2D).

Next, we set out to integrate these windows into segments with meaningful boundaries. Adjacent windows assigned to the same methylation state were merged into non-overlapping segments, allowing gaps of up to 1,000 bp between windows (Figure 1D). This gave 1,873,564 segments spanning 808,007,050 bp. By number, 54.7% of the segments were methylated, 41.8% transition and 3.4% unmethylated (Figure 2B); by length, they accounted for 72.2%, 23.6% and 4.2% of the segmented sequence, respectively (Supplementary Table S1). The number of segments per chromosome is shown in Supplementary Figure S3A.

**Figure 2.**
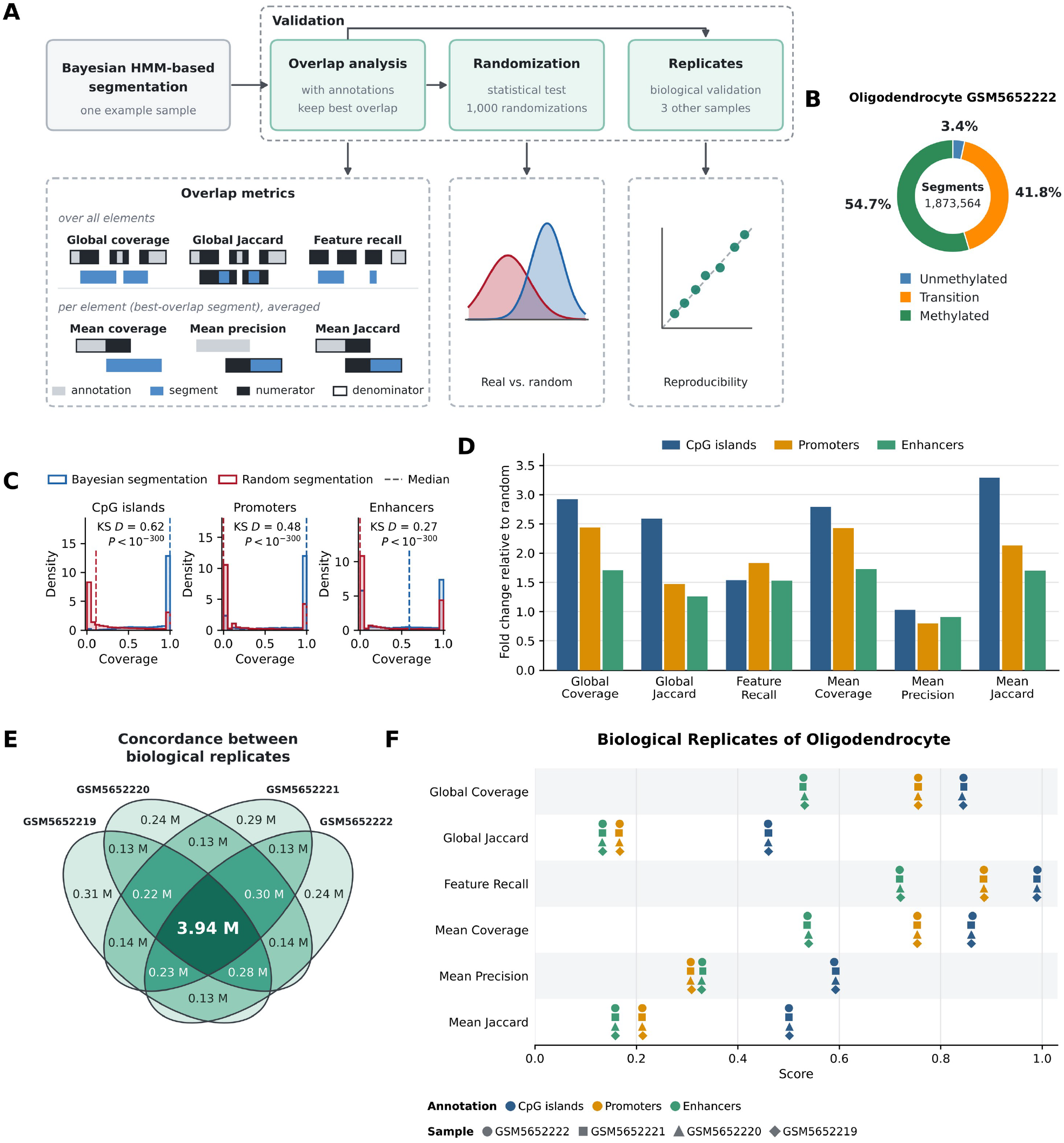
Bayesian segmentation of an oligodendrocyte methylome and its validation. (A) Validation workflow for one example sample (oligodendrocyte GSM5652222). The segmentation is evaluated by overlap with genomic annotations, by comparison with 1,000 randomized segmentations, and by replication in three other oligodendrocyte samples. Six overlap metrics are drawn schematically for annotation elements (grey) and segments (blue); dark fill marks the numerator and the outline the denominator. Global coverage, global Jaccard index and feature recall are computed over all elements; mean coverage, mean precision and mean Jaccard index are computed for each element and its best-overlapping segment and then averaged. (B) Composition of the 1,873,564 segments by methylation state (percentage of segments). (C) Coverage of CpG islands (n = 26,641), promoters (PLS; n = 45,890) and enhancers (pELS; n = 242,790) by the best-overlapping segment, for the Bayesian segmentation (blue) and one randomized segmentation (red). Dashed lines, medians. KS D, Kolmogorov–Smirnov statistic; P < 10^−300^for both the KS and the Mann–Whitney U test. (D) Fold change of each overlap metric relative to the mean of 1,000 random segments; a fold change of 1 is the random expectation. The empirical P-value is 0.001 for all metrics except mean precision for promoters and enhancers (P = 1). (E) Concordance between biological replicates. Venn diagram of the windows with the same chromosome, start position and methylation state in four oligodendrocyte samples; numbers are millions of windows, and the shading darkens with the number of samples that share a region. (F) Overlap-metric scores for CpG islands, promoters and enhancers in GSM5652222 (circles) and in three replicates: GSM5652221 (squares), GSM5652220 (triangles) and GSM5652219 (diamonds).

Segment lengths were strongly right-skewed, with many short segments and few long ones (Supplementary Figure S3B), and they differed among the states (Supplementary Table S1). Methylated and unmethylated segments had mean lengths of 569 bp and 523 bp (median 200 bp for both), whereas transition segments had a mean of 244 bp and a median of 50 bp, that is, a single window. Transition segments were therefore typically short intervals, consistent with their position between stable regions.

The segments spanned 28.9% of the WGBS-callable regions (808,007,050 of 2,796,121,515) and 28.0% of the autosomal reference genome (2,881,033,286 bp for chromosomes 1–22 of hg19; Supplementary Figure S3C). Coverage varied among chromosomes and was highest on chromosomes 19 and 17 (Supplementary Figure S3C). Because windows were joined across gaps of up to 1,000 bp, the segmented sequence is larger than the sequence of the modeled windows themselves (5,381,758 windows × 50 bp = 269 Mb, or 9.6% of the callable regions).

### 3.2 Evaluation of the segmentation output

To examine the genomic context of the segments, we intersected them with CpG islands and with ENCODE candidate cis-regulatory elements (cCREs), namely promoter-like (PLS) and proximal enhancer-like (pELS) signatures, and compared six overlap metrics with those of 1,000 random segments (Figure 2A; Sections 2.6–2.8).

The sets of random segments matched the real segments in terms of the number and length distribution of the segments on each chromosome and were placed without overlap within the callable regions (see Methods). The random expectation for global coverage is therefore close to the fraction of callable sequence spanned by the segments (about 30%).

Our real segments obtained from the single oligodendrocyte sample overlapped CpG islands, promoters and enhancers more significantly than random segments did. The distribution of per-element coverage was shifted toward complete coverage (Figure 2C; Kolmogorov–Smirnov D = 0.62, 0.48 and 0.27 for CpG islands, promoters and enhancers; P < 10^−300^). Global coverage was 0.845 versus 0.289 for randomized segmentations at CpG islands (2.92-fold), 0.756 versus 0.310 at promoters (2.44-fold) and 0.529 versus 0.310 at enhancers (1.71-fold) (Figure 2D; Supplementary Table S2). At least one segment overlapped 99.0% of CpG islands, 88.5% of promoters and 71.9% of enhancers (feature recall; random expectation 64.3%, 48.4% and 46.9%). Global Jaccard index, mean coverage and mean Jaccard index were also higher than random for the three classes (empirical P = 0.001, the minimum possible with 1,000 randomizations).

Mean precision, defined as the overlap between an element and its best-overlapping segment divided by the length of that segment, averaged over all elements, did not follow this pattern. It was close to random at CpG islands (fold change 1.03) and lower than random at promoters (0.80) and enhancers (0.91; empirical P = 1 for both). This indicates that the segments overlapping promoters and enhancers often extend beyond the element: the segmentation captures these elements but does not delimit their exact boundaries.

To assess reproducibility, the same workflow was applied to WGBS data from three other oligodendrocyte samples (GSM5652221, GSM5652220 and GSM5652219). The resulting segments were compared with the initial sample GSM5652222. Windows were nearly identical among the four samples: about 99% of the modeled windows were shared, and 5.33 million windows were retained in all four (Supplementary Figure S4A). Because windows lie on a fixed 50-bp grid, this mainly reflects common coverage filtering, and the informative question is whether shared windows receive the same state. When the same state assignment was also required, the agreement between pairs of samples was about 86%, and 3.94 million windows had the same state in all four samples (Figure 2E). Concordance depended on methylation state (Supplementary Figure S4B). Across the six sample pairs, 93–96% of unmethylated windows, 89–91% of methylated windows and 72–77% of transition windows kept their state. Differences involved adjacent states: transition windows changed mostly to methylated (21–27%), unmethylated and methylated windows changed to transition (4–7% and 9–11%, respectively), and direct switches between the unmethylated and methylated states were at most 0.2%.

Obtained segments across each of the 4 oligodendrocytes showed the same pattern. For the segments of the initial sample, 80–83% overlapped a segment of the same state in a replicate, 6– 7% overlapped a segment of an adjacent state, 0.01% overlapped a segment of the opposite state, and 11–13% overlapped no segment (Supplementary Figure S5A). The fractions with the same state were 91–94% for unmethylated, 84–87% for methylated and 67–71% for transition segments (Supplementary Figure S5B), so the states ranked as they did at the window level. The overlap metrics for all annotation classes were nearly identical among the four samples (differences ≤ 0.004; Figure 2F; Supplementary Table S3).

Together, these comparisons show that the windows, states, segments and annotation overlap of the initial sample were reproduced in three other oligodendrocyte samples, with transition regions being the least stable.

### 3.3 Construction of a segmentome atlas across human cell types

The reproducibility of our initial segments across oligodendrocyte biological replicates, together with the tissue-specific nature of DNA methylation, suggested that the segment boundaries might harbor cell-type specific information. To assess this, we applied our segmentation method to all 207 samples from the DNA methylation atlas previously published (Loyfer et al., 2023; 39 cell types; Figure 1E). In the resulting segmentome profiles, each sample is represented by its segments, with segment identity defined by chromosome, start and end position. Pooling the segments of all samples gave 27,095,738 unique segments, which formed a common feature space, and a binary presence matrix recorded whether each segment was present in each sample. Because identity depends on exact coordinates, segments that differ only slightly in their boundaries count as separate features. To reduce this redundancy, two segments were defined as representative of one another when their overlap was at least 90% of the length of the longer segment (see Methods). This relationship was found for 11,276,415 pairs of segments, involving 9,787,268 segments (36.1% of the pool). When one segment of a related pair was present in a sample, its counterpart was also recorded as present, which gave the “corrected presence matrix”.

The cumulative number of unique segments increased as sam-ples were added, but fewer new segments were added with each additional sample (Supplementary Figure S6). Later samples thus contributed progressively fewer previously unobserved segments, which suggests that the segment pool was approaching saturation for the cell types in this atlas. The plateau refers to the samples and cell types present in this cohort and does not mean that all human segments have been captured.

To test whether the corrected segmentome representation carries information about cell type identity, we retained the 28,000 most variable segments, embedded the samples with UMAP and clustered them by agglomerative hierarchical clustering (Euclidean distance, complete linkage), without using cell-type labels at any step (Figure 1F). The resulting dendrogram was then annotated with cell type labels (Figure 3).

**Figure 3.**
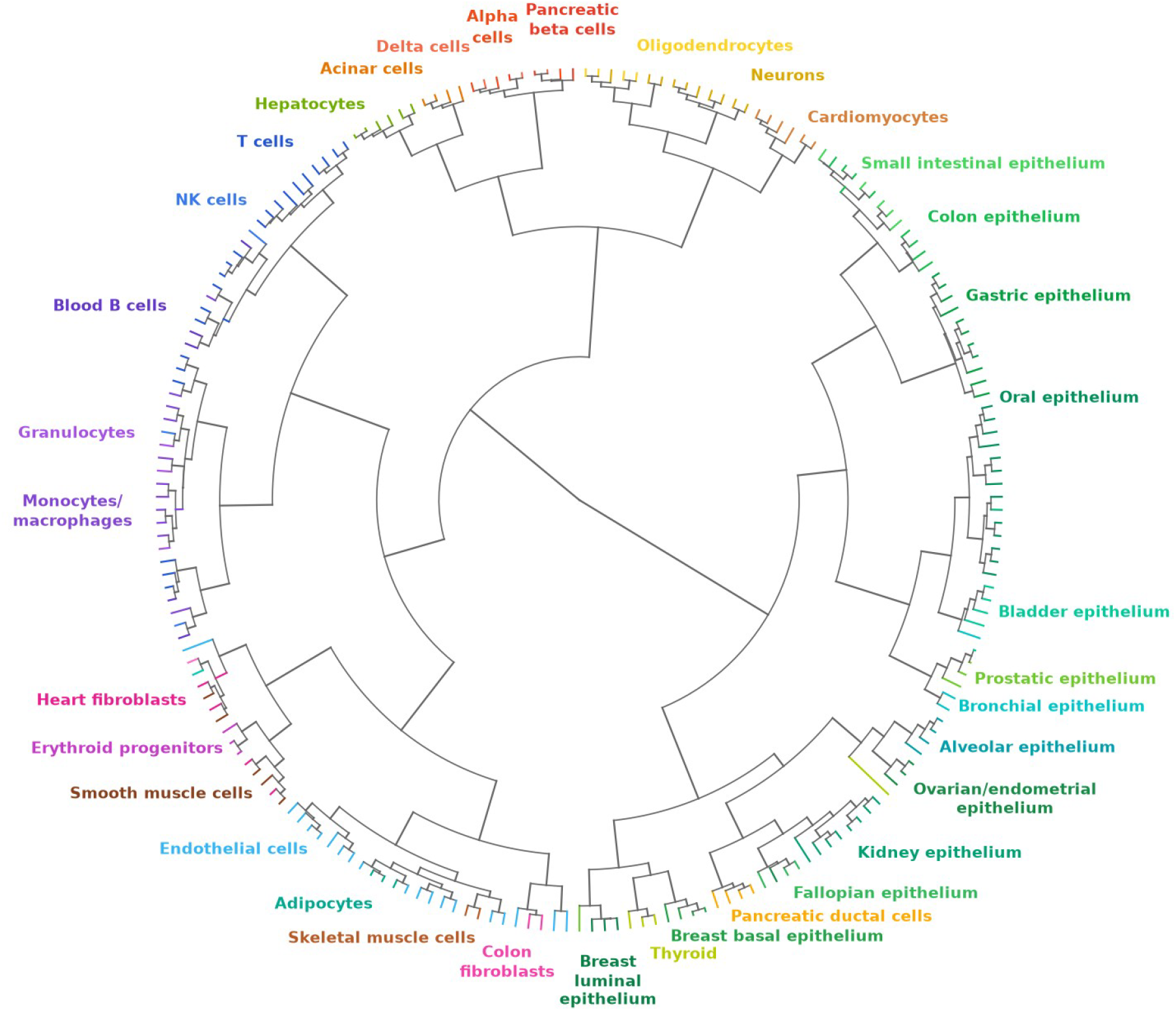
Hierarchical clustering of segmentome profiles by cell type. Branch colors and labels indicate the annotated cell type.

Samples of the same cell type generally clustered together, and related cell types formed larger branches. The blood cell types (T cells, NK cells, B cells, granulocytes and monocytes/macrophages) formed a single branch, and most epithelial cell types were found in a second large branch. Pancreatic alpha, beta, delta and acinar cells and hepatocytes were adjacent, as were neurons, oligodendrocytes and cardiomyocytes (Figure 3). Cutting the dendrogram into 39 clusters gave an adjusted Rand index of 0.55, an adjusted mutual information of 0.72 and a purity of 0.80; that is, 166 of the 207 samples were in a cluster whose majority cell type was their own.

Resolution was incomplete in some branches. Within the blood branch, samples of lymphoid and myeloid cell types were partly interleaved, and endothelial cells, adipocytes, colon fibroblasts and skeletal muscle cells shared one branch, whereas heart and colon fibroblasts were found in different branches. Overall, in an unsupervised analysis, the segmentome profiles exhibited information on cell type identity.

### 3.4 Identification of segmentome-derived cell-type-specific biomarkers with potential in liquid biopsy applications

Cell-type-specific segments were selected from the corrected presence matrix (target prevalence ≥ 70%, background prevalence < 2%, and ≤ 20% prevalence in any single non-target cell type), which gave 358,524 candidate segments across all 39 cell types (Figure 1G; Figure 4A). The top 25 candidates per cell type (968 segments in total) were used to compare the 39 cell types: the mean pairwise Euclidean distances computed from these seg-ments were low within a cell type and high between cell types (Figure 4B). Because the segments were selected for cell-type specificity in the same samples, this separation is expected and shows that the selected sets are specific.

**Figure 4.**
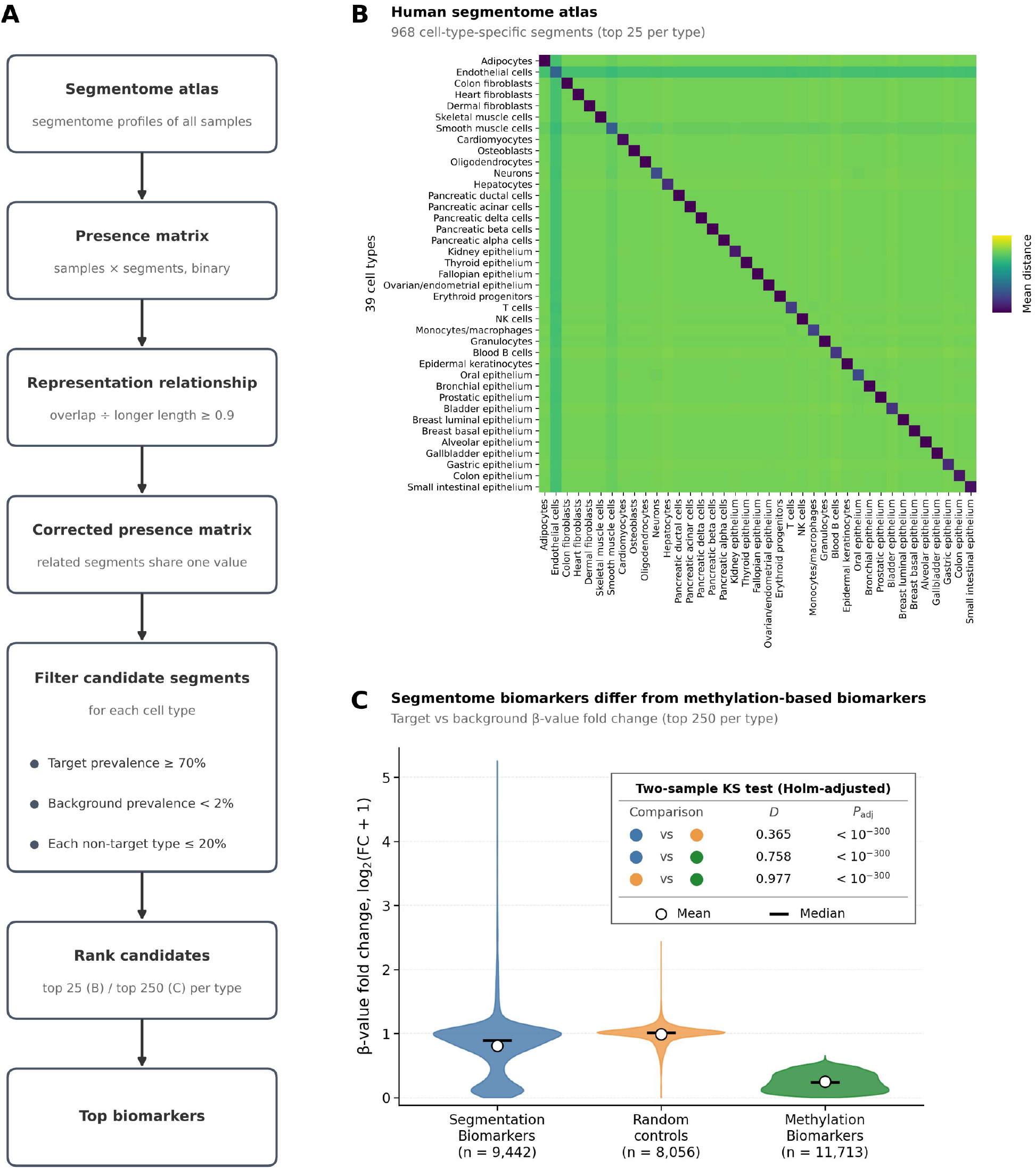
Identification and characterization of segmentome-derived cell-type-specific biomarkers. (A) Biomarker discovery workflow. Candidate segments were selected from the corrected presence matrix (target prevalence ≥ 70%, background prevalence < 2%, and ≤ 20% prevalence in any single non-target cell type) and ranked within each cell type; the top 25 (B) or top 250 (C) per cell type were retained. (B) Mean pairwise Euclidean distance between the 39 cell types, computed from the 968 segmentome biomarkers (top 25 per cell type). Low distances (dark) along the diagonal indicate similar profiles within a cell type. (C) Violin plots of the β-value fold change between target and background samples, shown as log2(fold change + 1), for segmentome biomarkers (n = 9,442), randomized control segments (n = 8,056) and the methylation biomarkers reported. (n = 11,713). Circles, mean; horizontal bars, median. The inset gives the two-sample Kolmogorov–Smirnov statistic (D) and the Holm-adjusted P value for each pair (P_adj < 10^−300^for all three).

Given the segmentome itself is defined on boundaries of DNA methylation, we next set out to test whether our segmentome-de-rived cell-type specific biomarkers are actually distinct from simple DMRs. To this end, we quantified the methylation levels of these biomarker segments. We calculated for each segment the β-value fold change between samples in the target (the cell type to which the segment was specific) and the background (all other cell types) classes and compared three sets (Figure 4C): the segmentome biomarkers (top 250 per cell type; n = 9,442), random control segments (n = 8,056; see Methods) and the methylation biomarkers identified in the original publication by Loyfer et al. (2023; n = 11,713).

The distributions of log_2_(fold change + 1) differed for all three pairs (two-sample Kolmogorov–Smirnov tests with Holm correction, P_adj < 10^−300^). The largest difference was between random controls and the methylation biomarkers (D = 0.977), followed by segmentome versus methylation biomarkers (D = 0.758) and segmentome biomarkers versus random controls (D = 0.365).

The methylation biomarkers had the lowest values, that is, the largest difference in methylation between target and background samples. The random controls were centered near 1 (no difference), and the segmentome biomarkers lay in between, with a broad distribution that overlapped with that of the controls (Figure 4C). The segmentome biomarkers therefore differ in methylation level between target and background samples much less than established methylation biomarkers do, consistent with their selection by segment presence rather than by methylation level difference. These results confirm that segmentome markers capture information that is distinct from that of conventional DMRs.

To evaluate our biomarkers on independent data, we generated segmentome profiles for 23 cell-free DNA (cfDNA) samples and their 23 paired white blood cell (WBC) samples (Loyfer et al., 2023). We next attempted to perform cell-type deconvolution analyses on these independent samples that contain mixed DNA fragments from multiple cell types. For this, we matched each mix-ture segmentome profile against the biomarker set identified on the previously described 39 cell types (see Methods; Figure 1H; Figure 5A).

**Figure 5.**
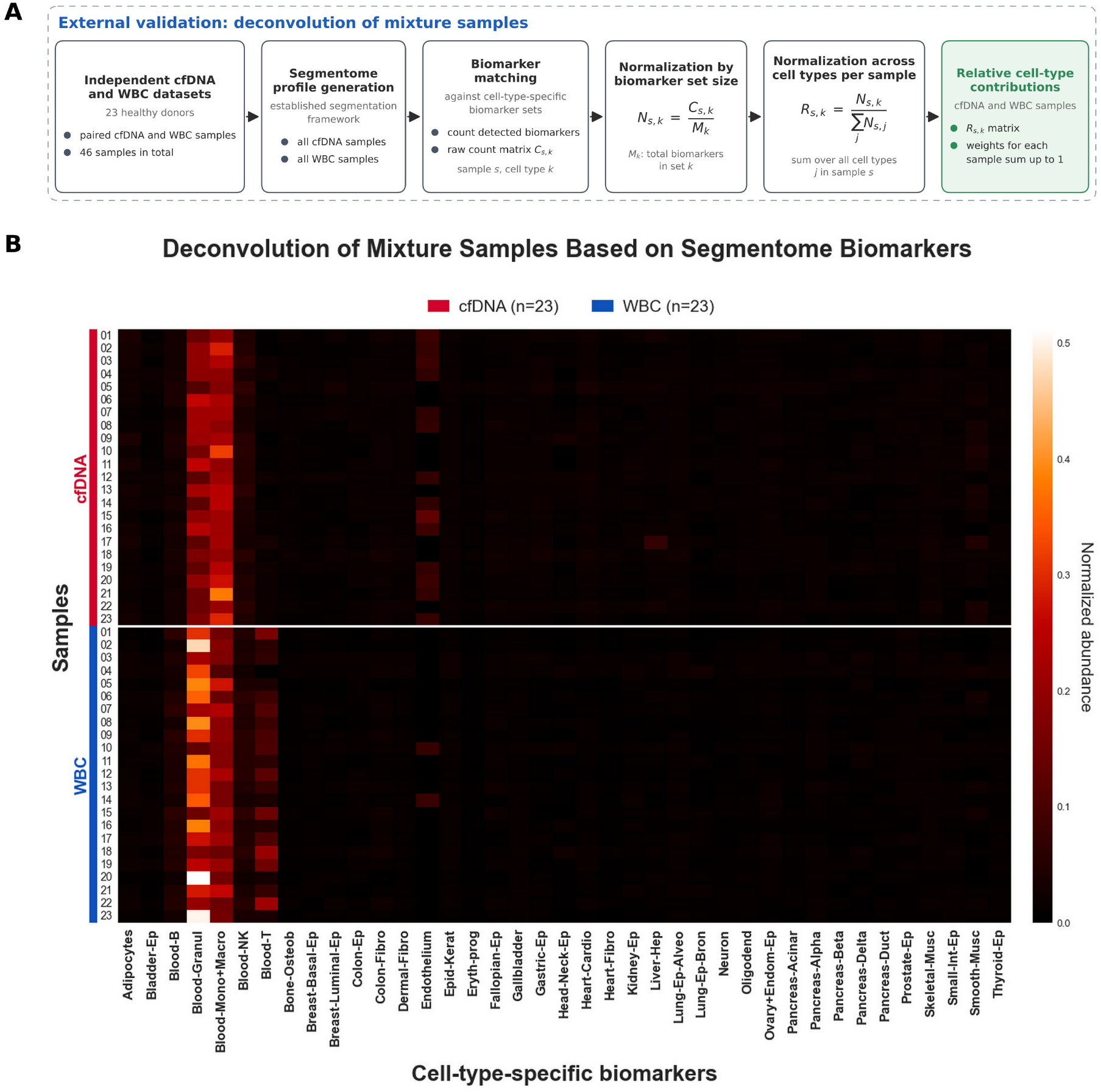
External validation of segmentome biomarkers by deconvolution of healthy cfDNA and WBC samples. (A) Workflow. Segmentome profiles were generated for 23 cell-free DNA (cfDNA) samples and their 23 paired white blood cell (WBC) samples. For each sample, the detected biomarkers of each cell type were counted by matching its profile to the full cell-type-specific biomarker sets (all biomarkers of each cell type; raw counts C), normalized by the number of biomarkers in the set (N = C/M) and then across cell types within the sample (R = N/ΣN), so that the values of each sample sum to one. (B) Heatmap of the normalized biomarker abundance (R) for the 39 cell-type-specific biomarker sets (columns) in cfDNA (red) and WBC (blue) samples (rows; rows 01–23 in each group are the same donors). Lighter colors indicate higher abundance.

The counts were normalized by the size of each biomarker set and then across cell types within each sample, so that the values of each sample sum to one (the values are relative biomarker abundances, not calibrated estimates of cell-type proportions, so the patterns should be read as comparative signatures rather than quantitative measurements of tissue contributions).

Blood cell types had the highest values in both sample types, but the patterns differed between WBC and cfDNA (Figure 5B). In cfDNA, the Blood-Monocyte+Macrophage set was the most prominent in most samples, followed by the Blood-Granulocyte set. In WBC samples, the Blood-Granulocyte set was the most prominent, and the Blood-T set (and, more weakly, the Blood-B set) was visible in WBC but not in cfDNA samples. A signal for the endothelial set appeared in some cfDNA samples and not in WBC samples. These patterns were consistent across donors, although the intensity of individual signals varied among samples. Non-blood sets had very low values in both sample types as expected.

These profiles show that segmentome biomarkers are able to delineate the cells of origin in cfDNA and WBC samples with multiple cell-type composition.

## 4 Discussion

The epigenome consists of multiple layers that are regulated in cells to undertake many important roles together in concert. These layers include DNA methylation, histone modifications, chromatin compaction, and non-coding RNA-assisted regulation.

Here we use the term regulome for the regulatory elements of the genome together with the epigenetic layers that control them. DNA methylation and the other layers are coupled in both directions. CpG islands are generally unmethylated, mark promoters and carry characteristic chromatin features (Deaton & Bird, 2011). Methylation is reduced at distal regulatory elements where transcription factors bind and chromatin is accessible (Stadler et al., 2011; Kundaje et al., 2015). The regulome is commonly annotated with HMMs that combine several histone marks into chromatin states (Ernst & Kellis, 2012). The segmentome is a model of the same kind, but it is inferred from the methylome alone. The overlap of our segments with CpG islands, promoters and enhancers is consistent with this view and suggests that part of the regulome can be inferred from methylation data. This has not yet been tested directly. The segments were compared with annotations that are not specific to the cell type, not with chromatin accessibility, histone modifications or three-dimensional contacts measured in the same cells.

When the DNA methylome is profiled using standard NGS-based methods at single base resolution, the main focus in data analysis is typically on identifying context-specifically hyperor hypomethylated regions and potentially drawing clinical applications from these. Here, we analyze this same data in a novel fashion to address a different question: how is methylation organized within a single cell type, and whether this organization can then be used to compare samples. We present the segmentome for the first time and show that methylome profiles can be leveraged to decipher more than CpG methylation patterns alone. Our results support this view in three ways. The inferred segments coincided with CpG islands and regulatory elements more than expected by chance; the segmentation of the initial sample was reproduced in other samples of the same cell type; and segmentome profiles retained information on cell identity across 39 cell types, although the model was developed on one oligodendrocyte sample and never saw cell-type labels. This evidence is indirect, however, and the paragraphs below discuss what each result does and does not show.

The most distinctive element of the model is the transition state. Transition windows were not simply intermediate in methylation level: their mean methylation was closer to that of methylated than of unmethylated windows, and they were distinguished by diverse read patterns and weak agreement between neighboring CpGs. In such regions the average methylation level is poorly informative, because the same average can arise from molecules that are uniformly intermediate or from a mixture of fully methylated and fully unmethylated ones. Within-sample heterogeneity scores were developed to capture this information and are complementary to the methylation level (Scherer et al., 2020). In many cancer types, disordered DNA methylation is reported as a cause or consequence of epigenetic instability (Hansen et al., 2011; Timp & Feinberg, 2013; Landau et al., 2014). The transition state can be seen as a segment-level counterpart of these measures. Its biological origin remains to be further resolved. Heterogeneous read patterns can reflect stochastic disorders, allele-specific methylation or a mixture of cell populations (Landau et al., 2014; Scherer et al., 2020), and sorted cell populations are not fully pure or homogeneous (Loyfer et al., 2023). The direction of transitions gives one clue. Transition windows switched into methylated windows much more often than to unmethylated windows, and direct transformations between the two stable states were rare, which suggests that transition windows lie mostly within or at the edges of methylated regions and less often next to unmethylated ones. Transition segments were also the least reproducible between replicates. This fits their heterogeneous nature, but it could equally reflect a less stable model fit in segments that are short by construction, and our data cannot distinguish between the two scenarios.

The segments most strongly overlapped with CpG islands, fol-lowed by promoters and enhancers. This agrees with the known link between unmethylated or lowly-methylated regions and regulatory activity (Deaton & Bird, 2011; Stadler et al., 2011). In MethylSeekR, unmethylated regions are mostly CpG islands, including promoters, and lowly-methylated regions corresponding to distal regulatory regions (Burger et al., 2013), and in the methylation atlas of Loyfer et al. (2023), cell-type-specific unmethylated regions were typically accessible and were proposed to be enhancers. The segmentome recovers this association without annotations, fixed methylation thresholds or labels, which suggests that the inferred states reflect organized methylation and not noise. Two features of the comparison probably explain why the association is weaker for enhancers. Enhancers are less CpG-dense than CpG islands and promoters (Stadler et al., 2011; Burger et al., 2013), so fewer of their windows can be modeled, and the cCRE registry is not specific to oligodendrocytes, so many registered elements need not be active, or unmethylated, in this sample. The low mean precision shows that the segments are coarser than the elements: they capture the regions in which promoters and enhancers lie but not their exact boundaries, as expected from the 50-bp windows and the merging of windows across gaps of up to 1,000 bp. The segmentome should therefore be read as a regional description. Neither enrichment nor reproducibility establishes function or positional accuracy.

The segmentome model introduced here differs from existing approaches in what it captures. DMR discovery methods locate differences between predefined groups, and the HMMs developed so far for methylation data mostly model these same differences (Wreczycka et al., 2017; Ji, 2019; Shen et al., 2017; Malonzo & Lähdesmäki, 2023). Methods that describe one methylome, such as MethylSeekR, rely on methylation level: unmethylated and lowmethylated regions are defined with a fixed methylation cut-off and a CpG-number criterion calibrated by a false discovery rate, and large partially methylated domains are found with a separate HMM (Burger et al., 2013). The segmentome model instead infers states without predefined thresholds, uses read-level features, and treats the transition between stable regions as a state of its own. The methylation atlas defined previously by Loyfer et al. (2023) is closer to our method in purpose. It segmented the genome into homogeneous blocks jointly across all samples by dynamic programming and selected markers as blocks that differ in methylation between cell types (Loyfer et al., 2023). Joint segmentation gives all samples common boundaries, which simplifies comparison, but the boundaries depend on the cohort. In our approach, the segmentome is computed for each sample independently with a fixed model, so a new sample can be segmented alone. The cost is that boundaries vary between samples, which produces a large and partly redundant pool of segments and made the representation relationship necessary. The two representations are therefore complementary, and a direct comparison with them, and with MethylSeekR, remains to be done.

Most notably, we show that the segmentome profiles retain in-formation on cell identity using a model that was completely agnostic to cell type labels. Related cell types were found in the most branches of clustering, as in the original atlas, where methylation grouped cell types by developmental origin (Loyfer et al., 2023). Because a segment is identified by its coordinates, the clustering reflects where boundaries occur, which depends on the local methylation landscape. The segmentome biomarkers were selected for their presence in one cell type and absence in the others, not for a methylation contrast, and their methylation differences between target and background samples were accordingly smaller than that of markers chosen for such a contrast. This is expected from the selection, and it indicates that segment presence reflects regulatory features of local organization that a mean methylation value does not show. It also means that these markers cannot be expected to match the sensitivities of markers chosen for maximal methylation contrast if methylation level alone is used as a feature.

Finally, as a proof of concept, we performed deconvolution analyses. In cfDNA and WBC samples, the signals of blood cell types dominated and differed between the two sample types, which is consistent with their different cell composition; granulocytes and monocytes/macrophages were also the main contributors in the fragment-level methylation-based deconvolution of healthy plasma by Loyfer et al. (2023).

Several limitations apply. The model’s resolution is fixed at 50 bp, and windows with fewer than two CpGs or without reads are not modeled, so the segmentome mainly describes the CpG-containing part of the genome. The validation relied on overlap with annotations, on replicates of one cell type, and on a null model that does not control for CpG density. Real segments can arise only in windows with enough CpGs, which are more frequent at CpG islands and promoters, so part of the enrichment may reflect where windows can be formed. Segment identity depends on exact coor-dinates, which makes the representation sensitive to small boundary differences that the representation relationship only partly corrects. Finally, the atlas consists of healthy sorted cells, the biomarkers were selected and evaluated in the same atlas, and the cfDNA analysis used 23 donors and a detection rule that requires an exact match, which is conservative.

Overall this study provides several useful insights. The partial independence of our cell-type specific segmentome markers from DMRs allows for the extraction of unique and complementary information from existing DNA methylome data. This may have potential to reduce the dependence on costly multiomic experiments for future studies by allowing for inference of some other regulome information from the technologies that produce DNA methylome only. If the segmentome proves robust, it could be used in several directions. Disordered methylation is a feature of disease (Landau et al., 2014), and heterogeneity scores can reveal diseaseassociated loci that average methylation does not (Scherer et al., 2020), so the transition state and the read-level features could be tested in tumors and in other conditions in which methylation heterogeneity changes. Segmentome biomarkers could complement methylation-contrast markers in cfDNA analysis once their signals are calibrated. Because methylation is depleted at or near domain boundaries in most lineages (Zhou et al., 2026), the transitions inferred here could also be related to the three-dimensional organization of the genome. These possibilities are prospective. They need independent cohorts, direct comparison with existing methods, and evaluation of reproducibility and clinical relevance.

## 5 Conclusion

In this study, we introduced for the first time, the *segmentome*, a segment-level representation of the DNA methylome that an unsupervised Bayesian HMM infers from WGBS data. The inferred segments are enriched at CpG islands and regulatory elements and shown to carry cell-type specific information across 39 human cell types. Here we showed potential applications in liquid biopsy contexts for tissue-of-origin inference. Further establishing the value of the segmentome in combination with other biomarkers will require additional external validation to demonstrate that it adds information beyond existing methods for DMR identification and methylation analyses.

