## Supplementary Figures for "The Segmentome: a novel representation of the DNA methylome identified by a Bayesian HMM framework"

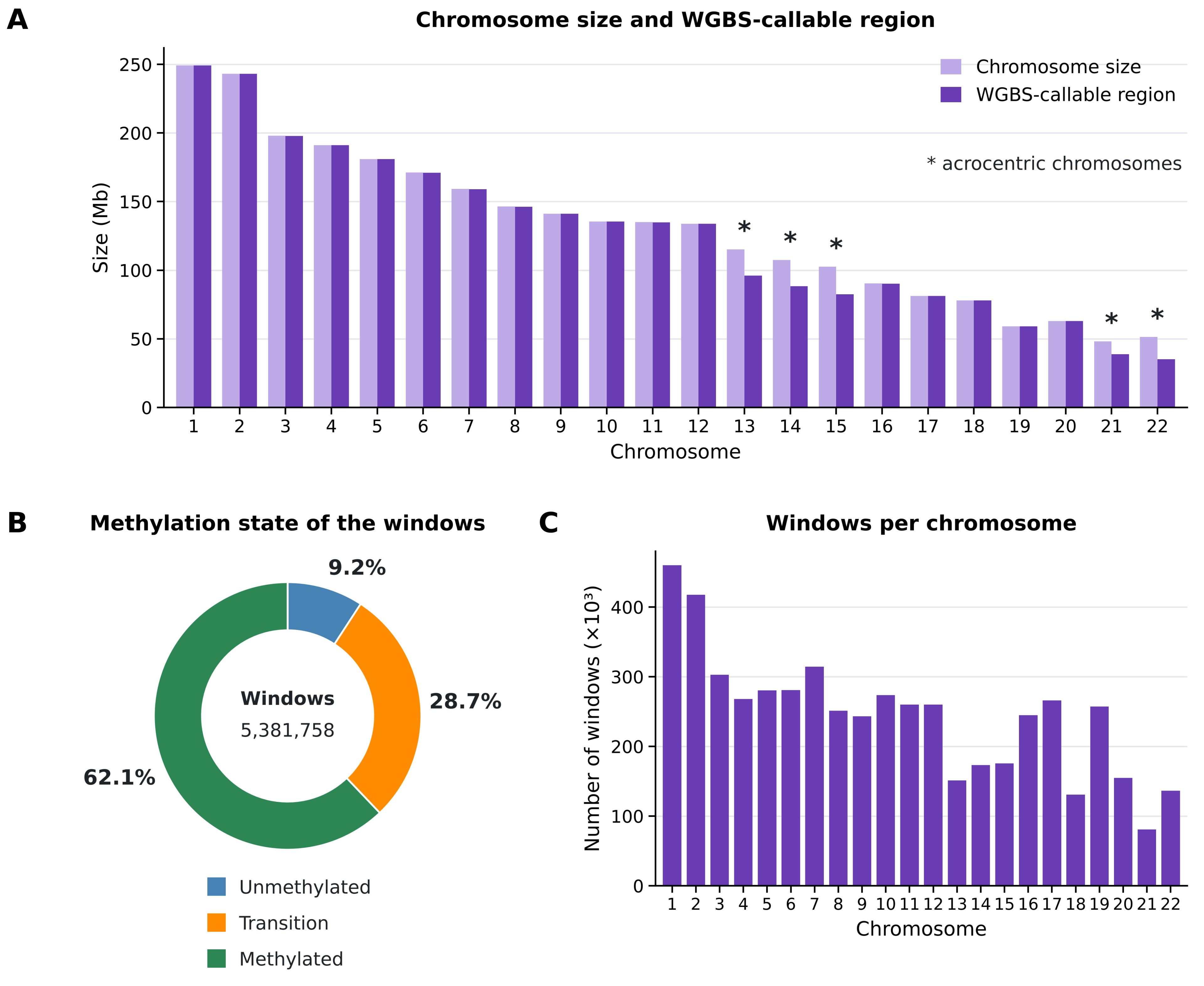
A
Chromosome size and WGBS-callable region
Chromosome size
WGBS-callable region
* acrocentric chromosomes
Size (Mb)
Chromosome
B
Methylation state of the windows
9.2%
28.7%
62.1%
Windows
5,381,758
Unmethylated
Transition
Methylated
C
Windows per chromosome
Number of windows (×10³)
Chromosome

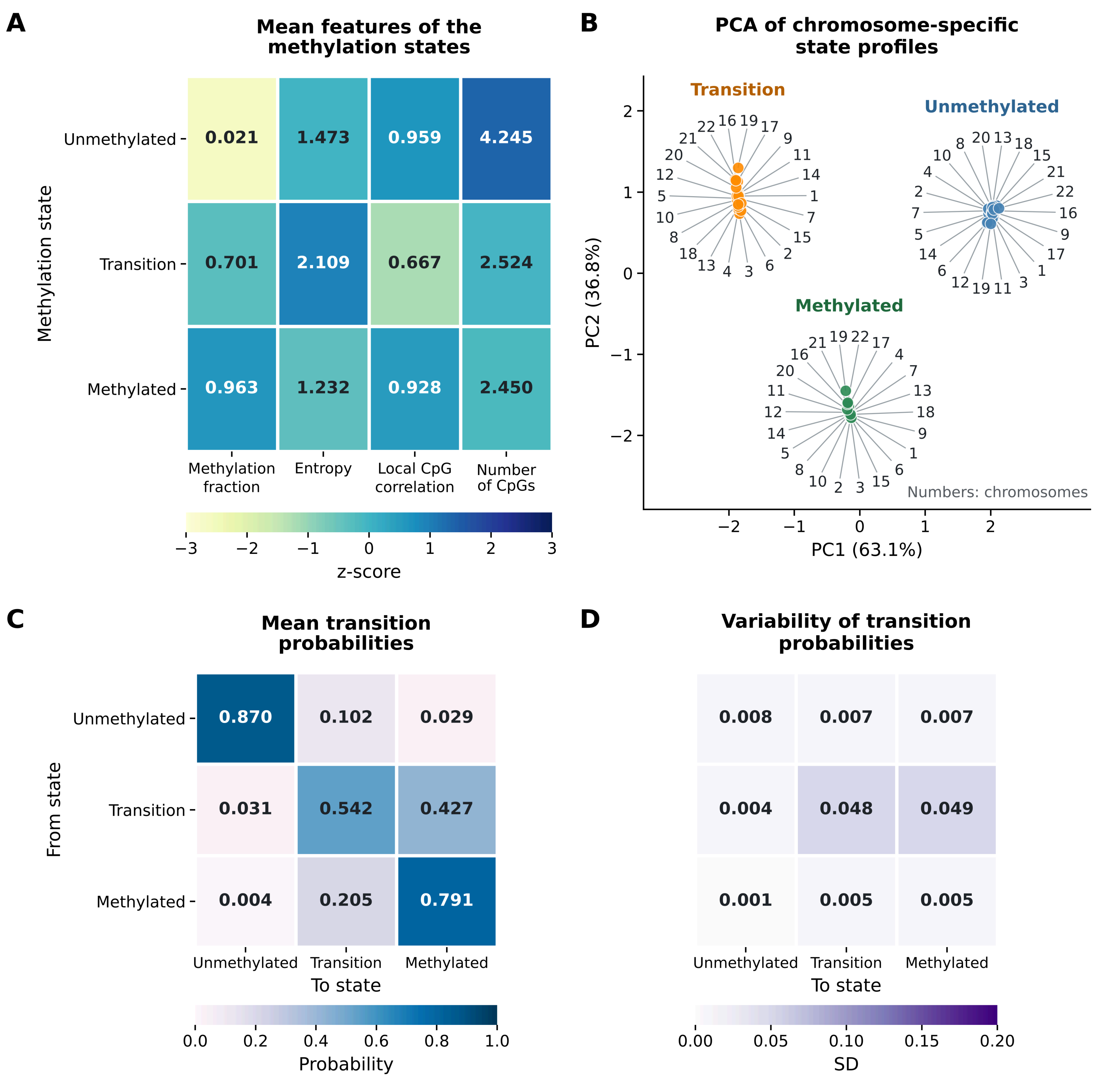
A
Mean features of the methylation states
Methylation state
Unmethylated
Transition
Methylated
0.021
1.473
0.959
4.245
0.701
2.109
0.667
2.524
0.963
1.232
0.928
2.450
Methylation fraction
Entropy
Local CpG correlation
Number of CpGs
−3
−2
−1
0
1
2
3
z-score
B
PCA of chromosome-specific state profiles
Transition
Unmethylated
Methylated
PC2 (36.8%)
PC1 (63.1%)
Numbers: chromosomes
C
Mean transition probabilities
From state
Unmethylated
Transition
Methylated
0.870
0.102
0.029
0.031
0.542
0.427
0.004
0.205
0.791
To state
Probability
0.0
0.2
0.4
0.6
0.8
1.0
D
Variability of transition probabilities
0.008
0.007
0.007
0.004
0.048
0.049
0.001
0.005
0.005
Unmethylated
Transition
Methylated
To state
0.00
0.05
0.10
0.15
0.20
SD

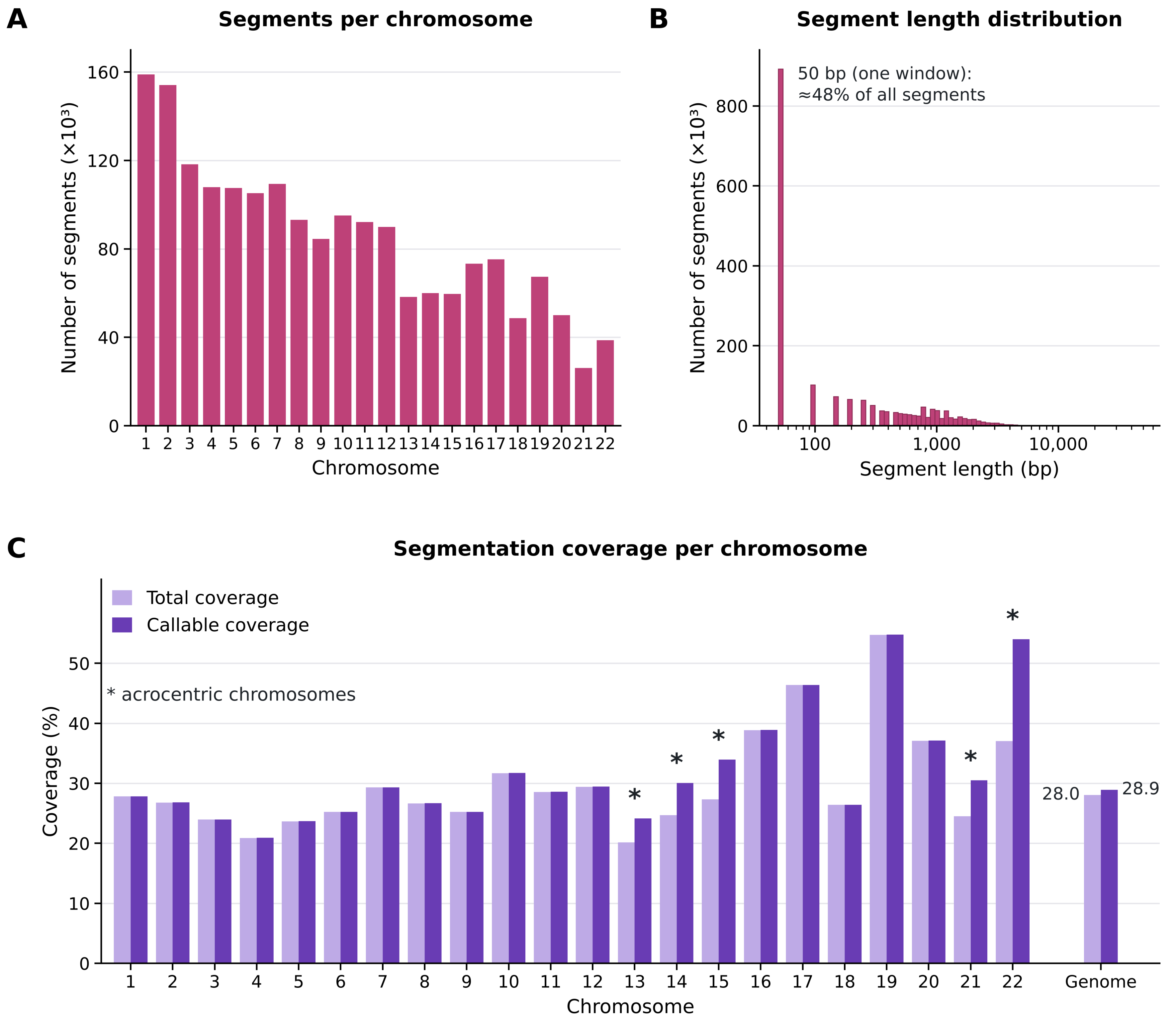
A
Segments per chromosome
Number of segments (×10³)
0
40
80
120
160
1 2 3 4 5 6 7 8 9 10 11 12 13 14 15 16 17 18 19 20 21 22
Chromosome
B
Segment length distribution
50 bp (one window):
≈48% of all segments
Number of segments (×10³)
0
200
400
600
800
100
1,000
10,000
Segment length (bp)
C
Segmentation coverage per chromosome
Total coverage
Callable coverage
* acrocentric chromosomes
Coverage (%)
0
10
20
30
40
50
*
*
*
*
*
28.0
28.9
1 2 3 4 5 6 7 8 9 10 11 12 13 14 15 16 17 18 19 20 21 22 Genome
Chromosome

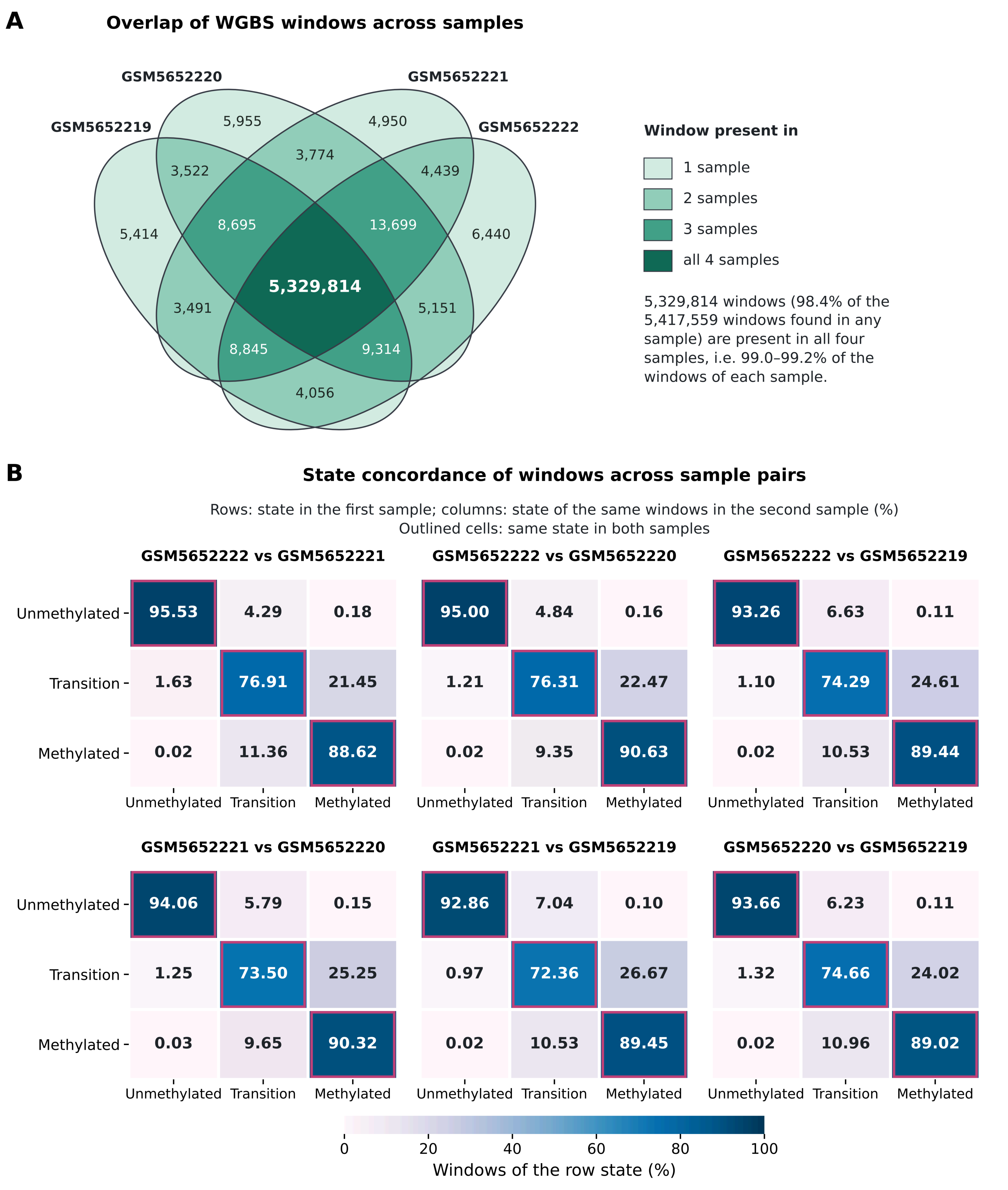
A
Overlap of WGBS windows across samples
GSM5652219
GSM5652220
GSM5652221
GSM5652222
5,955
4,950
3,522
3,774
4,439
5,414
8,695
13,699
6,440
5,329,814
3,491
5,151
8,845
9,314
4,056
Window present in
1 sample
2 samples
3 samples
all 4 samples
5,329,814 windows (98.4% of the 5,417,559 windows found in any sample) are present in all four samples, i.e. 99.0–99.2% of the windows of each sample.
B
State concordance of windows across sample pairs
Rows: state in the first sample; columns: state of the same windows in the second sample (%)
Outlined cells: same state in both samples
GSM5652222 vs GSM5652221
Unmethylated 95.53 4.29 0.18
Transition 1.63 76.91 21.45
Methylated 0.02 11.36 88.62
GSM5652222 vs GSM5652220
Unmethylated 95.00 4.84 0.16
Transition 1.21 76.31 22.47
Methylated 0.02 9.35 90.63
GSM5652222 vs GSM5652219
Unmethylated 93.26 6.63 0.11
Transition 1.10 74.29 24.61
Methylated 0.02 10.53 89.44
GSM5652221 vs GSM5652220
Unmethylated 94.06 5.79 0.15
Transition 1.25 73.50 25.25
Methylated 0.03 9.65 90.32
GSM5652221 vs GSM5652219
Unmethylated 92.86 7.04 0.10
Transition 0.97 72.36 26.67
Methylated 0.02 10.53 89.45
GSM5652220 vs GSM5652219
Unmethylated 93.66 6.23 0.11
Transition 1.32 74.66 24.02
Methylated 0.02 10.96 89.02
Unmethylated Transition Methylated
0 20 40 60 80 100
Windows of the row state (%)

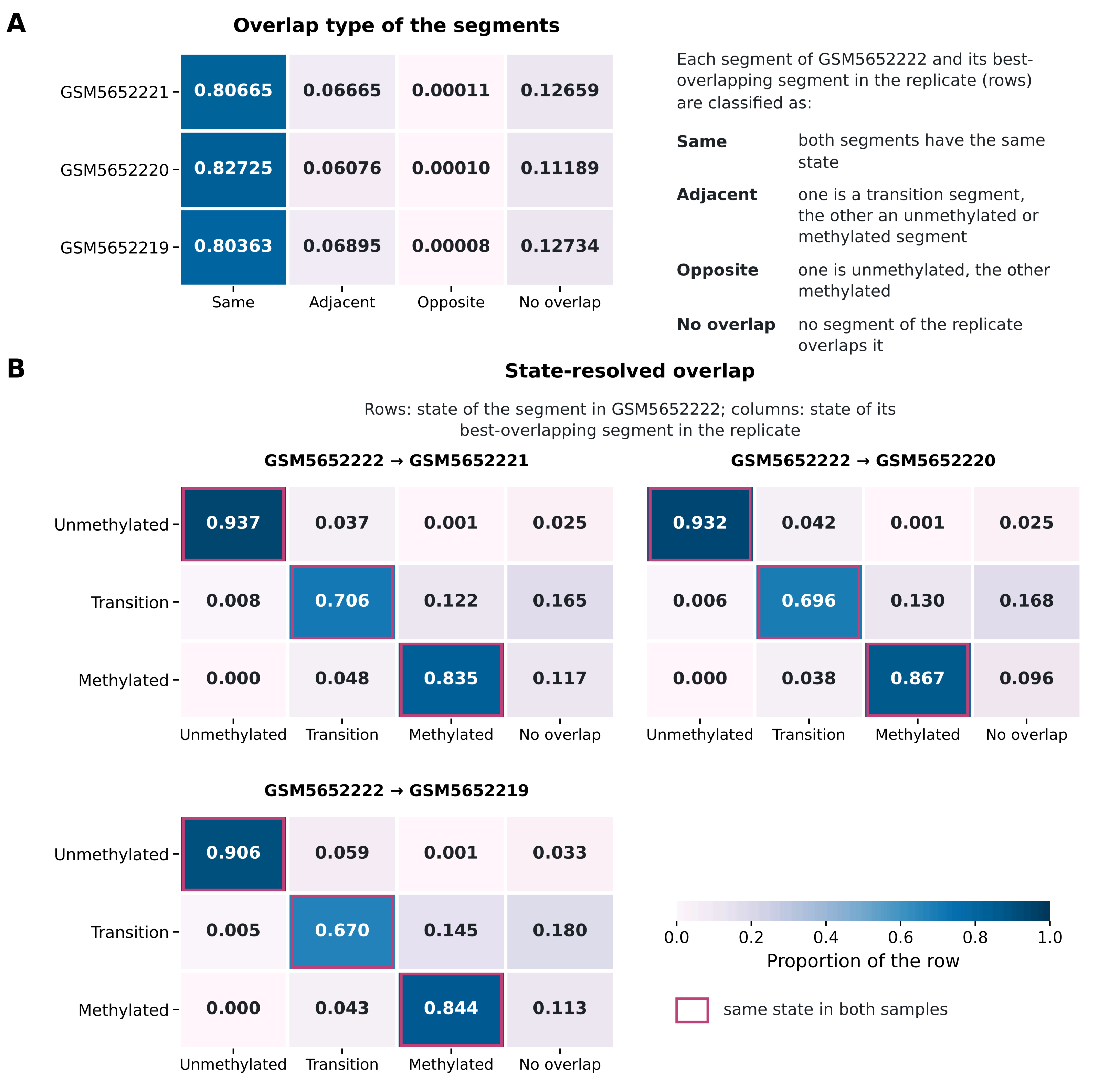
A
Overlap type of the segments
Same
Adjacent
Opposite
No overlap
GSM5652221
0.80665
0.06665
0.00011
0.12659
GSM5652220
0.82725
0.06076
0.00010
0.11189
GSM5652219
0.80363
0.06895
0.00008
0.12734
Each segment of GSM5652222 and its best-overlapping segment in the replicate (rows) are classified as:
Same
both segments have the same state
Adjacent
one is a transition segment, the other an unmethylated or methylated segment
Opposite
one is unmethylated, the other methylated
No overlap
no segment of the replicate overlaps it
B
State-resolved overlap
Rows: state of the segment in GSM5652222; columns: state of its best-overlapping segment in the replicate
GSM5652222 → GSM5652221
Unmethylated
Transition
Methylated
No overlap
Unmethylated
0.937
0.037
0.001
0.025
Transition
0.008
0.706
0.122
0.165
Methylated
0.000
0.048
0.835
0.117
GSM5652222 → GSM5652220
0.932
0.042
0.001
0.025
0.006
0.696
0.130
0.168
0.000
0.038
0.867
0.096
GSM5652222 → GSM5652219
0.906
0.059
0.001
0.033
0.005
0.670
0.145
0.180
0.000
0.043
0.844
0.113
0.0
0.2
0.4
0.6
0.8
1.0
Proportion of the row
same state in both samples

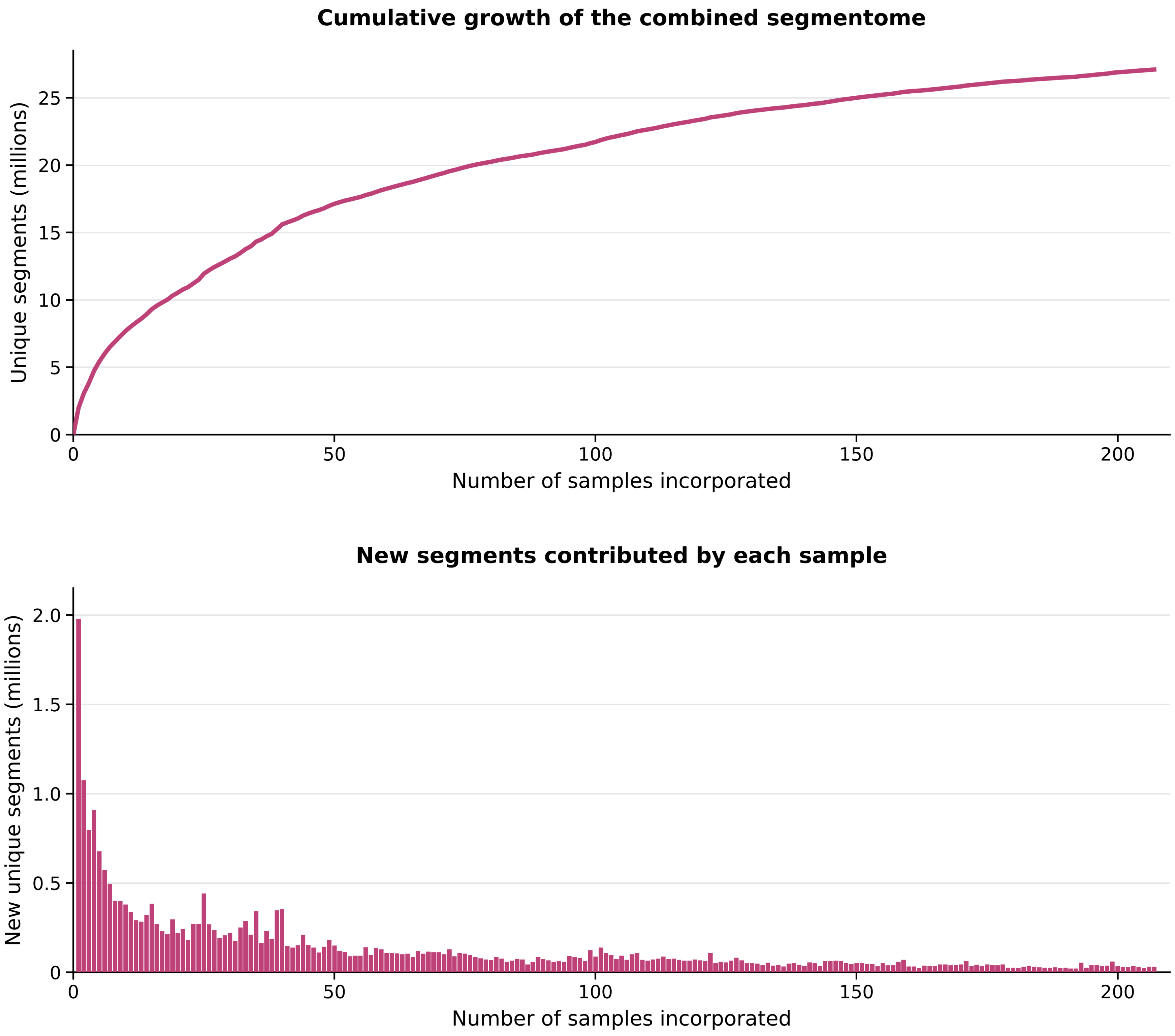
Cumulative growth of the combined segmentome
Unique segments (millions)
0
5
10
15
20
25
0
50
100
150
200
Number of samples incorporated
New segments contributed by each sample
New unique segments (millions)
0
0.5
1.0
1.5
2.0
0
50
100
150
200
Number of samples incorporated
